# Foliar gas exchange, morphology, and cannabinoid contents of three hemp varieties in southwest Texas

**DOI:** 10.1101/2025.03.09.642247

**Authors:** V. S. John Sunoj, Xuejun Dong, Madhumita V. Joshi, Russell W. Jessup, Daniel I. Leskovar, David D. Baltensperger

## Abstract

In the US, a high level (≥ 0.3%) of intoxicating Δ-9-tetrahydrocannabinol (THC) threatens farm-scale production of industrial of Hemp (*Cannabis sativa* L. ssp. *sativa*), but the linkage between THC and major physiol-morphological traits of hemp is not well-known. This study aims to characterize the variations in physiological and/or morphological parameters and cannabinoid contents of three hemp varieties, i.e., Berry Blossom, Painted Lady, and Skipper. Diurnal foliar gas exchange, chlorophyll fluorescence, water potential, and canopy temperature were measured on five clear days in the 2022 growing season, and cannabinoids were measured at peak flowering using high-performance liquid chromatography. Allometric equations were developed to use easily measured biomass or morphological variables to predict variables that are more difficult to measure. The diurnal foliar gas exchange of the three hemp varieties was largely unaffected by the high temperatures of southwest Texas, with Berry Blossom and Skipper showing the highest and lowest photosynthesis, respectively, and Painted Lady having the most efficient stomatal control of gas exchange. Although the rooting depth of Berry Blossom was shallower than that of the two other varieties, there was no evidence showing the effect of rooting habit on the physiology of the studied hemp varieties, which was presumably due to the lack of water stress in our experiment. Nor were there significant differences in the cannabinoid contents in relation to environmental and varietal responses, as the measured THC contents of all three varieties were under 0.3%. Overall, the three hemp varieties showed different behavior strategies in southwest Texas.

## 1 INTRODUCTION

Hemp (*Cannabis sativa* L. ssp. *sativa*) is an industrial multifaceted annual crop cultivated to produce fiber, grain, oil, and biologically active secondary metabolites such as cannabinoids, including medically important cannabidiol (CBD), and terpenes (Fike, 2016; Lazarjani et al., 2020). For decades, hemp had been banned in several countries due to its relationship with marijuana (*Cannabis sativa* L.), with levels of the intoxicating cannabinoid Δ-9-tetrahydrocannabinol (THC) above 0.3% (Ehrensing, 1998; USDA, 2000; Williams, 2020; Zhang et al., 2021). Recently, many countries began to legalize the production of hemp as an agricultural commodity. The allowable maximum level for total THC varies across nations, with the United States, Canada, China, and the EU defining < 0.3% as legal threshold (DeCarlo and Weaver, 2023), which indicates that THC concentration determines the fate of hemp cultivation and its products (Cherney and Small, 2016; Jiang et al., 2021; Toth et al., 2021). However, at the state level in the U.S., the allowable THC threshold varies, although most of the states legalize cannabis use in at least some form. For example, in Kansas, the THC cap for cannabis flower is 0%, but it is set at 30% and 35% in Mississippi and Tennessee, respectively. Twelve other states, including Texas, only allow cannabis products with low THC (https://www.oberk.com/marijuanalawsbystate).

Variations in abiotic factors, e.g., the amount of incident light, photoperiod, soil nutrients and water intake, groundwater availability, and day/night temperatures, can lead to changes in plant photosynthetic and/or other physiological traits. In hemp, these traits include flowering time, sex characteristics, biomass accumulation, seed nutritional and oil compositions (fatty acid profile), yield (grain and fiber) and yield quality (Hall et al., 2012; Tang et al., 2017; Herppich et al., 2020; Zhang et al., 2021; Sunoj V. S. et al., 2023). Most importantly, these variations can change the cannabinoid profile, particularly, the percentages of THC and CBD (Schluttenhofer and Yuan, 2017; Petit et al., 2021; Payment and Cvetkovska, 2023). Numerous studies have been conducted in hemp to understand the influence of different environmental factors on photosynthesis, e.g., photoperiod (Gajdošik et al., 2022), light intensity (Rodriguez-Morrison et al., 2021), heavy metal contamination (Linger et al., 2005; Shi et al., 2009; Sun et al., 2022), soil fertility (Tang et al., 2017; De Prato et al., 2022b; Saloner and Bernstein, 2022), microbes (De Prato et al., 2022a; Sun et al., 2022), light spectra (Islam et al., 2021; Jenkins, 2021; Cheng et al., 2022), water deficit (Tang et al., 2018; Caplan et al., 2019; Herppich et al., 2020; Jiang et al., 2021; Gill et al., 2022), and high temperatures (Chandra et al., 2011; Herppich et al., 2020). The production and presence of cannabinoids as secondary metabolites in hemp, for example, were frequently assumed to play a certain role in plant’s stress protection (Latta and Eaton, 1975; Pate, 1999; Flores-Sanchez and Verpoorte, 2008). However, data collected from open-field conditions where the hemp plants were directly interacting with the natural environment, was relatively limited (Herppich et al., 2020), and the key questions of the relationships between cannabinoid concentration and plant stress responses, as has been comprehensively reviewed by Payment and Cvetkovska (2023), still remain unanswered with site-specific data support.

Transpirational cooling associated with the deep-root traits plays a significant role in mitigating the effect of heat stress in crops such as wheat and chickpea (Pinto and Reynolds, 2015; Purushothaman et al., 2015; Deva et al., 2020). Hemp’s deep root trait and high root biomass have been cited as suitable for sustainable, low-input cropping systems (Amaducci et al., 2008; Gill et al., 2023). In subtropical species, plant rooting depth was observed to respond to increased temperature (Luo et al., 2020). However, we are unaware of the direct data that supports on whether a higher root depth/biomass in hemp can help ameliorate heat stress through increased water uptake, and if so, whether the moderation of heat stress is in turn associated with modified cannabinoid production. These questions are especially pertinent for regions prone to high temperature stress. The current study was conducted in southwest Texas to understand the potential interactions of the photosynthetic traits (gas exchange and chlorophyll fluorescence), morphological traits (shoot/root biomass), and the cannabinoid traits (CBD and THC) in hemp and to demonstrate a better strategy to screen hemp varieties for region-specific adaptations. We posed the following questions: (a) Were photosynthetic parameters of the studied hemp varieties significantly affected by the naturally occurring high temperature in southwest Texas? (b) Were hemp’s deep-root traits able to significantly reduce the canopy temperatures during the high-temperature hours of selected clear days, and did they have any influence on photosynthetic responses and leaf water potential? (c) Were there any salient trends of the cannabinoid profile being strongly associated with the high-temperature responses of the hemp varieties? (d) Were there any traits—photosynthetic, morphological, or cannabinoid in nature—that demonstrate the adaptability of the studied hemp genotypes to the climatic conditions of southwest Texas?

## 2 MATERIALS AND METHODS

### 2.1 Field site, plant material, and crop management

The experiment was conducted from April to September 2022 in an open field at Texas A&M AgriLife Research and Extension Center at Uvalde, Texas, USA (latitude 29°12′34”N, longitude 99°45′03”W; elevation 276 m). The field site has a semiarid climate (Figure 1) with clay soil (Fine-silty, mixed, active, Hyperthermic Aridic Calciustolls) of the Uvalde series (USDA, 1970). Three hemp varieties, i.e., Berry Blossom, Painted Lady, and Skipper (Davis Farms of Oregon, LLC, OR, USA) that yielded relatively high biomass among 16 varieties in a 2021 test at the same site (Uvalde) were selected to study the genotypic variations in physiological characteristics and cannabinoid profile. In April 2022, these three varieties were tested in Uvalde.

**FIGURE 1.**
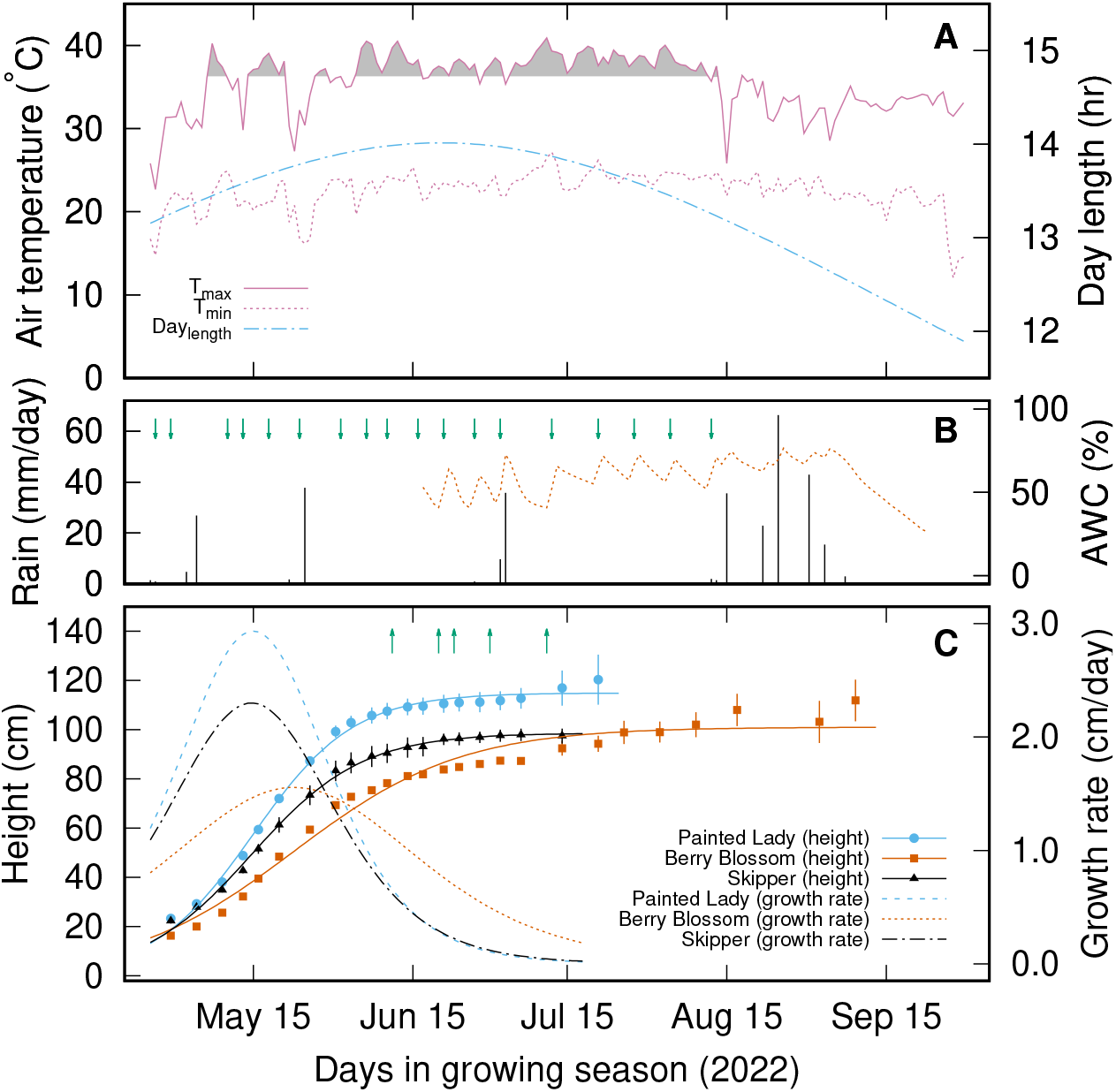
Environmental conditions and growth characteristics of three hemp genotypes used in this study at Uvalde in 2022. Daily maximum and minimum temperatures and day length (panel **A**), and daily rainfall (panel **B**) were measured from April 25 to September 30, 2022 at a weather station located at the field site. Daily available water capacity (AWC) in the top 60 cm of soil profile (expressed as percentage of the total available soil water, shown by the dashed line in panel **B**) was estimated based on the volumetric water content measured at the 15 cm, 30 cm and 60 cm soil depths. The data values of AWC from April 29 to June 16 were lost as a result of an operational mistake. The shaded area in panel **A** indicates the days when the daily maximum temperature was greater than 36.3 °C (temperature at which photosynthetic rate was estimated to be maximum, as seen Fig. 6). In panel **C**, data of the measured plant heights are expressed as symbols (error bars indicate ± one standard errors of the means with *n* = 5). A logistic model (Mead et al., 2003) was used to fit the height growth data of three hemp genotypes measured twice a week from late April to late August (**C**). The dashed lines represent the height growth rates of the there hemp genotypes, which were obtained by taking the first derivatives of the obtained best-fit curves (shown as solid lines in **C**). The down arrows in panel **B** indicate the days when irrigation was given through drip, which maintained the available soil water capacity around 63% in the top 60-cm of soil profile during the major growth periods. The up arrows in panel **C** indicate the days, i.e., June 11 (47 DAT), June 20 (56 DAT), June 23 (59 DAT), June 30 (66 DAT), and July 11 (77 DAT), when the diurnal leaf gas exchange rates, canopy temperatures, and leaf water potentials were measured for the three hemp genotypes. DAT: days after transplanting.

The total area of the Uvalde field site was 0.077 ha. Before planting, a composite soil sample was collected and analyzed (Soil, Water, and Forage Testing Laboratory, Texas A&M University, College Station, Texas, USA). The recommended rate of nitrogen (113 kg N ha^−1^) was applied to the field site in the form of urea granules before transplanting hemp seedlings and irrigation was provided through a drip system. During the experiment period, the drip system maintained a total available soil water capacity around 63% in the top 60 cm of soil profile along with the rainfall received (313 mm from April 25 to September 16, 2022; see Figure 1B). Throughout the experimental period, soil water content was measured using a set of capacitive soil moisture sensors (Model GS3), and data were recorded using an EM-50 data logger (Decagon Devices Inc [METER group], Washington, USA). Sensors were installed at 15.2 cm, 30.5 cm, and 61 cm soil depths and the daily average values of soil moisture contents (recorded in the datalogger once an hour; see Supplementary Figure S1) were used for estimating the total available soil water capacity, defined as the percentage of total available soil water to plants in the top 60-cm of soil profile. The soil water contents at the field capacity and wilting point for the Uvalde clay soil were estimated to be 0.35 and 0.18, respectively, based on prior work (Raman, 2018). To visualize the changes of the soil water status, a set of Watermark sensors were also installed and connected to an AM-400 datalogger (Mike Hansen Company, East Wenatchee, Washington, USA; also see Supplementary Figure S1). Routine weather data were recorded in a weather station located at the Research Center.

Twenty-nine-day-old hemp seedlings grown in pot trays were transplanted to the raised beds (1 m wide × 24 m long) on April 25, 2022. The experimental design was a randomized complete block design (RCBD). There were five blocks, each with four plants per variety used for this study. The spacing between plants was 1 m, with 2 m between blocks. Sunflower plants were grown at the border of each block (raised beds on both sides of the hemp block) as a wind barrier with a plant-to-plant spacing of 30-40 cm. Twenty-one days after transplanting (DAT), post-emergent herbicide Fusilade DX (Fluazifop-P-butyl; Syngenta Crop Protection AG; Basel, Switzerland) was applied. Further, at the time of the appearance of initial symptoms of leaf miner (after 9 DAT) and fall armyworm (*Spodoptera frugiperda*; after 61 DAT), biological insecticides Monterey LG6135 (Monterey Lawn and Garden Products Inc, California, USA) and DiPel Pro DF (Valent Bioscience, Illinois, USA), respectively, were applied, which helped prevent the hemp plants from insect attacks. Three to four male plants, once identified at early flowering, were removed promptly from the field plots. Weed control was done by hand.

### 2.2 Diurnal foliar gas exchange, chlorophyll fluorescence, canopy temperature, and water potential

Foliar gas exchange rates, such as net photosynthesis rate (*A*), stomatal conductance (*g*_*s*_), and intercellular CO_2_ concentration (*C*_*i*_) were measured using a portable photosynthesis system (Li-6400XT; LI-COR, Lincoln, Nebraska, United States). The instantaneous water use efficiency and carboxylation efficiency were calculated from the ratio of *A* to *E* and *A* to *C*_*i*_, respectively (Rymbai et al., 2014). The measurement was made on physiologically mature leaves from three phenotypically similar plants of each of the three varieties (see Supplementary Figure S1B). Due to the distances among the five blocks (separated by sunflower borders), the foliar gas exchange measurement was only made on plants in one of the five blocks. Five clear days from June 11 to July 11, 2022 (i.e., 47, 56, 59, 66, and 77 DAT; see the up arrows in Figure 1C) were selected to make the gas exchange and related physiological measurements (see see Supplementary Figure S2 for the hourly weather data of the five days). On each day, a minimum of five measurements were recorded from the middle part of the leaves of each selected plant at five different times (8:00 am, 10:00 am, 12:00 pm, 2:00 pm, and 4:00 pm). The CO_2_ concentration of the leaf chamber interior was set to 400 *µ*mol mol^−1^ with the use of a 6400-01 CO_2_ mixer installed with a pure CO_2_ cylinder (12 g). The system flow rate was set to 500 *µ*mol s^−1^ and the leaf fan was operating at high speed. Simultaneously, prior to the gas exchange measurement and at each time interval of a specific DAT, ambient light intensity incident on plant leaves was measured using an external quantum sensor attached to the leaf chamber and the same intensity of photosynthetically active radiation (PAR) was provided to the leaf inside the leaf chamber with a 6400-02B LED light source. The measurements were recorded under ambient relative humidity and temperature, and with a fixed leaf area of 2 × 3 = 6 cm^2^ exposed to the chamber cuvette. The sensor head of the photosynthesis system was placed in shade during the waiting time between measurement intervals.

In each cycle of foliar gas exchange measurement, leaf chlorophyll fluorescence transients, canopy temperature (*T*_*C*_), and leaf water potential (*ψ*_*L*_) were also measured. A minimum of four chlorophyll fluorescence and three *ψ*_*L*_ measurements were recorded from each variety at each measurement interval. Maximum photochemical efficiency of photosystem II (PSII; *F*_*v*_ /*F*_*m*_) and thylakoid membrane damage (*F*_0_/*F*_*m*_) were measured from fully opened mature leaves after 30 min of dark adaptation using a chlorophyll fluorometer (OS1p; OptiSciences, Hudson, New Hampshire, USA). The *F*_*v*_ /*F*_*m*_ was calculated as *F*_*v*_ /*F*_*m*_ = (*F*_*m*_ − *F*_0_)/*F*_*m*_ ; where *F*_0_ and *F*_*m*_ are the minimum and maximum fluorescence, respectively, in the dark-adapted state. A light pulse intensity of 3000 *µ*mol m^−2^ s^−1^ with a duration of 3 sec was applied to the leaf to generate *F*_*m*_.

Leaf water potential (*ψ*_*L*_) was measured using a pressure chamber instrument (Model 615; PMS Instrument Company, Albany, Oregon, USA), for which the high pressure was provided by compressed nitrogen gas. Canopy temperature (*T*_*C*_) was measured using an infrared radiometer with a handheld meter (Model MI-210, Apogee Instruments Inc, Logan, Utah, USA).

### 2.3 Cannabinoid profiling

For cannabinoid profiling, inflorescence samples were collected from the selected plants at BBCH (Biologische Bundesantalt, Bundessortenamt and CHemische Industrie, Germany) stage 65-67 (Mishchenko et al., 2017), which was characterized by abundant formation of capitate trichomes and fattening of the inflorescence buds. Samples were air-dried in an air-conditioned room (21-25 °C) for 6-8 weeks and ground into a fine powder using a Shardor coffee grinder. Approximately 500 mg of the ground material was mixed with 4 ml HPLC (high-performance liquid chromatography) grade methanol in a 20 mL vial, which was ultrasonicated at 20°C for 10 min using an ultrasonic water bath to extract cannabinoid. The extracted solution was then diluted twice, and the concentrations of 16 cannabinoids (CBC, CBCA, CBD, CBDA, CBDV, CBDVA, CBG, CBGA, CBL, CBN, CBNA, THCA, THCD8, THCD9, THCV, THCVA) were quantified using an Agilent 1220 Infinity Series HPLC with an autosampler installed. The complete package of sample preparation, HPLC calibration and automated measurement system (Cann-ID) as developed by the Ionization Labs (Austin, Texas, USA) was followed in this work. To adjust for the average moisture content of the air-dried samples, the quantities of the total CBD and total THC were calculated as (CBDA × 0.877) + CBD, and (THCA × 0.877) + THCD9, respectively.

### 2.4 Plant height, biomass accumulation, and root traits

Throughout the experiment period, plant height was measured twice a week from the root crown to the tip of the inflorescence of the main stem using a measuring stick (precision 1 mm). The plant height data were used to calculate the plant growth curve and height growth rate (Figure 1C). The total above-ground biomass accumulation was calculated by cutting the plants from the base and drying them at 70°C in a forced-air oven until a constant weight was obtained. To obtain the roots after cutting the plants, soil samples in the immediate vicinity of the plants were collected using a 10-cm diameter soil auger considering the root crown as the center point. For each plant, ten soil samples within the top 1-m depth profile were collected, each was 10 cm deep and 10 cm in diameter, except the first sample near the root crown region where the soil volume collected was 10 cm deep and 25 cm in diameter (done in order to quantify the horizontal distribution of roots). Six hundred soil samples (10 samples/plant × 4 plants/plot × 5 plots/variety × 3 varieties = 600 samples) were collected, and each of them was promptly put in a 3.78 L ziploc bag and stored in a −20°C freezer, awaiting further processing. All the soil samples were processed in small batches within a period of 5 months. Two days before processing a specific batch of the samples, frozen soil bags were thawed slowly by keeping them in a cold room at approximately 6°C.

After thawing, the soil was transferred to a tub, and roots are gently washed and filtered through sieves with different mesh sizes (top mesh [2 mm] and lower mesh [0.5 mm];U.S.A. Standard Testing Sieves, Milwaukee, Wisconsin, USA). Root washing was repeated until no root appeared after filtering. Washed roots were collected in plastic bottles/vials filled with 20% ethanol solution. For root samples from the 0-10 cm and 10-20 cm depth intervals, large bottles (300 cm^3^) were used to accommodate the typically larges sample sizes. Before collecting the roots from the root crown, the diameter of the root crown was measured using a digital caliper (CD-6” CS, Mitutoyo Group, Japan). If thick roots (diameter > 2.0 mm) were found in a sample, they were carefully separated from the remaining fine roots (diameter < 2.0 mm) and placed in a coin envelop and oven-dried at 70°C. Later the mass of the thick roots were added to calculate the total root mass at specific soil depth for a specific plant. Due to the typically large root sizes at the top two layers (0-10 and 10-20 cm depth intervals), only a sub-sample of fine roots from each of the samples of these layers was used in root length measurement. Specifically, the cleaned roots were floated in a water-filled tray to minimize cluttering and overlapping of rootlets, and scanned into computer to create a digital image, which was then analyzed using the WinRHIZO software ver. 2020 (Regent Instruments Inc., Canada) to calculate total root length (cm). The root length density of each of the soil samples (volume ≈ 4906 cm^3^ for the depth interval of 0-10 cm, and 1130 cm^3^ for the remaining depth intervals) was expressed as cm cm^−3^.

After scanning, all the fine roots from each sample were separated from the water using a funnel lined with a piece of filter paper (quantitative, 415; 15 cm; VWR International, USA). The filter paper together with roots was kept in a forced-air oven at 70°C until a constant mass was obtained. Using an analytical balance (VWR-160AC.N, VWR International, LLC), the mass of the filter paper was subtracted from that of fine roots plus filter paper combined to obtain the root dry mass.

### 2.5 Statistical analysis

The experimental design for the field study was a randomized complete block design (RCBD). There were three hemp varieties and five blocks (four plants per variety per block) used in this study. Three replications (*n* = 3) per variety from a single block were used for recording all the physiological traits. At the same time, five replications (*n* = 5) per variety from five blocks were used for recording all the morphological traits. Data on physiological traits (diurnal foliar gas exchange, chlorophyll fluorescence, leaf water potential, and canopy temperature), and morphological traits (plant height, biomass accumulation, and root traits) of the three varieties were analysed using Analysis of Variance (ANOVA) or Analysis of Means (ANOM), and the multiple comparisons of were made using the Tukey method in Minitab (Minitab LLC. Ver.21, State College, Pennsylvania, USA) with *α* = 0.05. The quantitative relationships of the major gas exchange variables were also analyzed by linear/nonlinear regression or visualized using the contour plot method available in Minitab.

A unified stomatal model (USM) of Medlyn et al. (2011) was fitted to the measured diurnal course data of leaf gas exchange in order to detect potential differences in stomatal behavior among hemp varieties. The model approximates the values of stomatal conductance as:

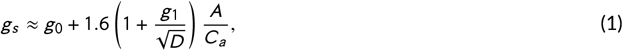

where *A* is net photosynthetic rate (*µ*mol m^−2^s^−1^), *C*_*a*_ is the CO_2_ concentration at the leaf surface (*µ*mol mol^−1^), *D* is the leaf-to-air vapor pressure deficit (kPa), *g*_0_ is minimum stomatal conductance (mol m^−2^s^−1^) and *g*_1_ is proportional to the marginal water cost of carbon gain 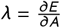 as used by Cowan and Farquhar (1977) : 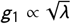 (also see Hari et al. (1986)). Since the magnitude of the intercept parameter *g*_0_ of Eqn. 1 approximately equals zero (Medlyn et al., 2011), we chose to omit it when fitting the USM to the measured gas exchange data, as seen in other studies (Duursma et al., 2019; Gardner et al., 2023). The nonlinear regression procedure (the Gauss-Newton method) available in Minitab was used to do the model fitting. Since the model has three independent variables (i.e., *D, A* and *C*_*a*_), we chose to pool all the gas exchange data measured across different days from a particular hemp variety in order to estimate the value of parameter *g*_1_ of Eqn. 1 corresponding to that variety. The statistical differences in model parameter (*g*_1_) among hemp varieties were compared using ANOVA with GraphPad Prism 6 (Version 6.07 for Windows, 2015, GraphPad Software, San Diego, California, USA), along with the Tukey’s multiple comparisons test (*α* = 0.05). To visualize the stomatal behavior described by Eqn. 1, *g*_*s*_ was regressed against 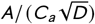 and the differences in the regression slope of different hemp varieties were compared using ANCOVA with Prism 6.

The allometric relationships between pairs of major biomass variables, including shoot fresh/dry mass, root fresh/dry mass, and inflorescence fresh/dry mass, were established using linear regressions on log-transformed data according to Troughton (1956). Data values in support of all the figures were graphed and visualized using gnuplot (Williams and Kelley, 2019).

## 3 RESULTS

### 3.1 Diurnal variation in physiological traits of different hemp varieties

As seen in Table 1, the effects of time of day (*T*), days after planting (*D*), and the interaction of them (*T* × *D*) on the eleven physiological traits were highly significant (*P* < 0.0005), while only some of the effects involving variety (*V*) were significant. This suggests that the effects involving *V* are more interesting, and we will focus more on them in reporting the results. While there was no significant varietal effect on leaf water potential (*ψ*_*L*_), maximum photochemical efficiency of PSII (*F*_*v*_ /*F*_*m*_), and thylakoid membrane damage (*F*_0_/*F*_*m*_), there were some effects in photosynthesis (*A*), carboxylation efficiency (*A*/*C*_*i*_), stomatal conductance (*g*_*s*_), transpiration (*E*) and instantaneous water use efficiency (*A*/*E*). These mainly reflect the trend that the values of these variables for Skipper were significantly lower than those of Berry Blossom or Painted Lady.

**TABLE 1.**
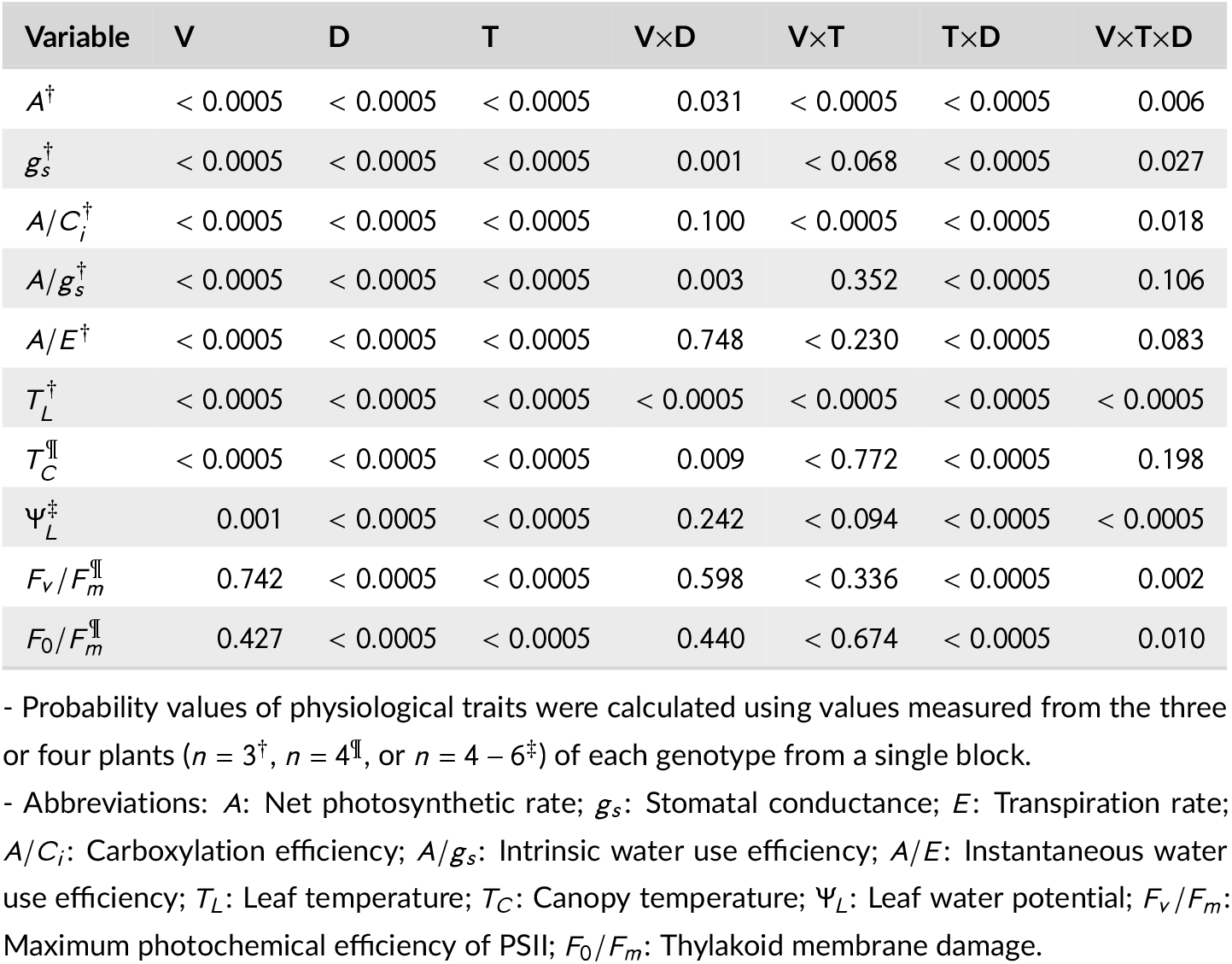
Probability values for variety (V), days after transplanting (D), time of day (T), and interactions (V×D, V×T, T×D, and V×T×D) effects on physiological parameters of three hemp varieties in Uvalde in 2022.

*V* × *D interactions*. As seen in Table 2, photosynthesis (*A*) of Skipper was significantly lower than that of two other varieties on 47, 56, and 77 DAT, while on two other days (59 and 66 DAT), it was only lower than that of Berry Blossom. The significant effect of *V* × *D* on *g*_*s*_ (*P* = 0.001 in Table 1) was due to the significantly higher value of Berry Blossom than that of Skipper on 47 and 59 DAT, but not on three other days (see Table 2). The strong *V* × *D* interaction effect on *E* was detected (*P* < 0.0005), because transpiration of Painted Lady was significantly lower than two other varieties only on 59 DAT, while on four other days Painted Lady and Skipper had similar transpiration, being significantly lower than that of Berry Blossom.

**TABLE 2.** Differences in daytime means in selected physiological trait variables of three hemp varieties over 5 clear days in Uvalde in 2022. Results are based on Tukey pairwise comparisons for the interaction of variety (V) and days after transplanting (D) as shown in Table 1.

| Variable | Variety | 47 DAT | 56 DAT | 59 DAT | 66 DAT | 77 DAT |
| --- | --- | --- | --- | --- | --- | --- |
| $A$ ($\mu\text{mol m}^{-2}\text{s}^{-1}$) | Painted Lady | <sup>a</sup>20.39 | <sup>a</sup>12.96 | <sup>ab</sup>20.45 | <sup>ab</sup>16.26 | <sup>ab</sup>12.00 |
|  | Berry Blossom | <sup>a</sup>22.64 | <sup>a</sup>13.44 | <sup>a</sup>23.15 | <sup>a</sup>18.79 | <sup>a</sup>13.59 |
|  | Skipper | <sup>b</sup>15.06 | <sup>c</sup>8.57 | <sup>bc</sup>19.43 | <sup>bc</sup>14.65 | <sup>c</sup>8.01 |
| $A/C_i$ ($\text{mol m}^{-2}\text{s}^{-1}$) | Painted Lady | <sup>ab</sup>0.069 | <sup>ab</sup>0.051 | <sup>a</sup>0.075 | <sup>ab</sup>0.063 | <sup>ab</sup>0.050 |
|  | Berry Blossom | <sup>a</sup>0.076 | <sup>a</sup>0.056 | <sup>a</sup>0.083 | <sup>a</sup>0.071 | <sup>a</sup>0.055 |
|  | Skipper | <sup>c</sup>0.050 | <sup>bc</sup>0.038 | <sup>a</sup>0.071 | <sup>bc</sup>0.056 | <sup>c</sup>0.033 |
| $g_s$ ($\text{mol m}^{-2}\text{s}^{-1}$) | Painted Lady | <sup>ab</sup>0.69 | <sup>a</sup>0.024 | <sup>ab</sup>0.56 | <sup>a</sup>0.32 | <sup>a</sup>0.18 |
|  | Berry Blossom | <sup>a</sup>0.86 | <sup>a</sup>0.20 | <sup>a</sup>0.71 | <sup>a</sup>0.42 | <sup>a</sup>0.24 |
|  | Skipper | <sup>c</sup>0.48 | <sup>a</sup>0.11 | <sup>bc</sup>0.53 | <sup>a</sup>0.30 | <sup>a</sup>0.12 |
| $A/g_s$ ($\mu\text{mol}^{-1}$) | Painted Lady | <sup>a</sup>33.44 | <sup>b</sup>69.80 | <sup>a</sup>46.78 | <sup>a</sup>59.06 | <sup>a</sup>67.13 |
|  | Berry Blossom | <sup>a</sup>29.67 | <sup>b</sup>72.18 | <sup>a</sup>36.33 | <sup>a</sup>50.58 | <sup>a</sup>58.87 |
|  | Skipper | <sup>a</sup>38.34 | <sup>a</sup>86.00 | <sup>a</sup>42.84 | <sup>a</sup>56.75 | <sup>a</sup>64.36 |
| $E$ ($\text{mmol m}^{-2}\text{s}^{-1}$) | Painted Lady | <sup>b</sup>8.02 | <sup>ab</sup>3.98 | <sup>b</sup>8.99 | <sup>bc</sup>7.56 | <sup>b</sup>7.52 |
|  | Berry Blossom | <sup>a</sup>10.05 | <sup>a</sup>5.16 | <sup>a</sup>12.43 | <sup>a</sup>10.11 | <sup>a</sup>10.40 |
|  | Skipper | <sup>b</sup>6.85 | <sup>bc</sup>3.17 | <sup>a</sup>11.36 | <sup>ab</sup>8.25 | <sup>b</sup>6.62 |
| $A/E$ ($\mu\text{mmol}^{-1}$) | Painted Lady | <sup>a</sup>2.98 | <sup>a</sup>3.22 | <sup>a</sup>2.30 | <sup>a</sup>2.14 | <sup>a</sup>1.75 |
|  | Berry Blossom | <sup>b</sup>2.52 | <sup>b</sup>2.75 | <sup>ab</sup>1.94 | <sup>ab</sup>1.89 | <sup>ab</sup>1.48 |
|  | Skipper | <sup>b</sup>2.37 | <sup>b</sup>2.70 | <sup>bc</sup>1.76 | <sup>bc</sup>1.79 | <sup>bc</sup>1.34 |
| $T_L$ ($^{\circ}\text{C}$) | Painted Lady | <sup>a</sup>37.91 | <sup>c</sup>35.31 | <sup>c</sup>36.21 | <sup>bc</sup>36.26 | <sup>c</sup>40.68 |
|  | Berry Blossom | <sup>a</sup>38.33 | <sup>b</sup>36.82 | <sup>b</sup>37.18 | <sup>ab</sup>36.97 | <sup>b</sup>41.94 |
|  | Skipper | <sup>a</sup>38.57 | <sup>a</sup>37.80 | <sup>a</sup>38.04 | <sup>a</sup>37.72 | <sup>a</sup>43.17 |
| $T_C$ ($^{\circ}\text{C}$) | Painted Lady | <sup>a</sup>33.02 | <sup>a</sup>33.33 | <sup>a</sup>31.92 | <sup>a</sup>32.58 | <sup>a</sup>37.32 |
|  | Berry Blossom | <sup>b</sup>31.21 | <sup>a</sup>33.12 | <sup>a</sup>31.10 | <sup>a</sup>32.43 | <sup>a</sup>37.39 |
|  | Skipper | <sup>a</sup>32.62 | <sup>a</sup>33.40 | <sup>a</sup>31.29 | <sup>a</sup>32.65 | <sup>a</sup>38.06 |

- For a specific variable, different lowercase letters indicate significant difference among/between hemp varieties ( $P < 0.05$ ).
- Abbreviations: $A$ : Net photosynthetic rate; $g_s$ : Stomatal conductance; $E$ : Transpiration rate; $A/C_i$ : Carboxylation efficiency; $A/g_s$ : Intrinsic water use efficiency; $A/E$ : Instantaneous water use efficiency; $T_L$ : Leaf temperature; $T_C$ : Canopy temperature.

*V* × *T interactions*. Diurnal values of *A, A*/*C*_*i*_, *g*_*s*_, *A*/*E* and *A*/*g*_*s*_ were more variable than those of leaf temperature (*T*_*L*_), canopy temperature (*T*_*C*_), *F*_0_/*F*_*m*_ and *ψ*_*L*_ (Figures 2 and 3). Detailed results on the interaction effects of variety and time of day on physiological traits are shown in Supplementary Tables S1-S5. Several significant effects in *A, E, A*/*C*_*i*_ and *T*_*L*_ are seen by referring to column four of Table 1 and Figures 2-3. The significant *V* × *T* interaction effect on *A* mainly reflected in the lower value of *A* in Skipper, as well as the reduction of *A* in Painted Lady in the afternoon hours (*P* < 0.05), while in the morning hours, both Painted Lady and Berry Blossom had higher values of *A* than Skipper (Table 1; Figure 2A-E). The significant *V* × *T* interaction effect on *A*/*C*_*i*_ largely mirrored the trend seen in *A* (Table 1; Figure 2F-J). The afternoon reduction of *A* in Painted Lady was also accompanied with the reduction in *E* in this variety, especially on 47, 59 and 77 DAT (Figure 2U-Y). This effect, as well as the trends that the value of *E* in Berry Blossom was the highest and that of Skipper the lowest among the three varieties, resulted in the significant *V* × *T* interaction effect in *E* (*P* < 0.0005). Finally, at different times of a day, Skipper tended to have the warmest leaf temperature (*T*_*L*_), and Pained Lady the coolest, among the three varieties (Figure 3P-T). For example *T*_*L*_ of Painted Lady in the morning from 8:00 am to 12:00 pm was 2.3°C lower than that of Skipper.

**FIGURE 2.**
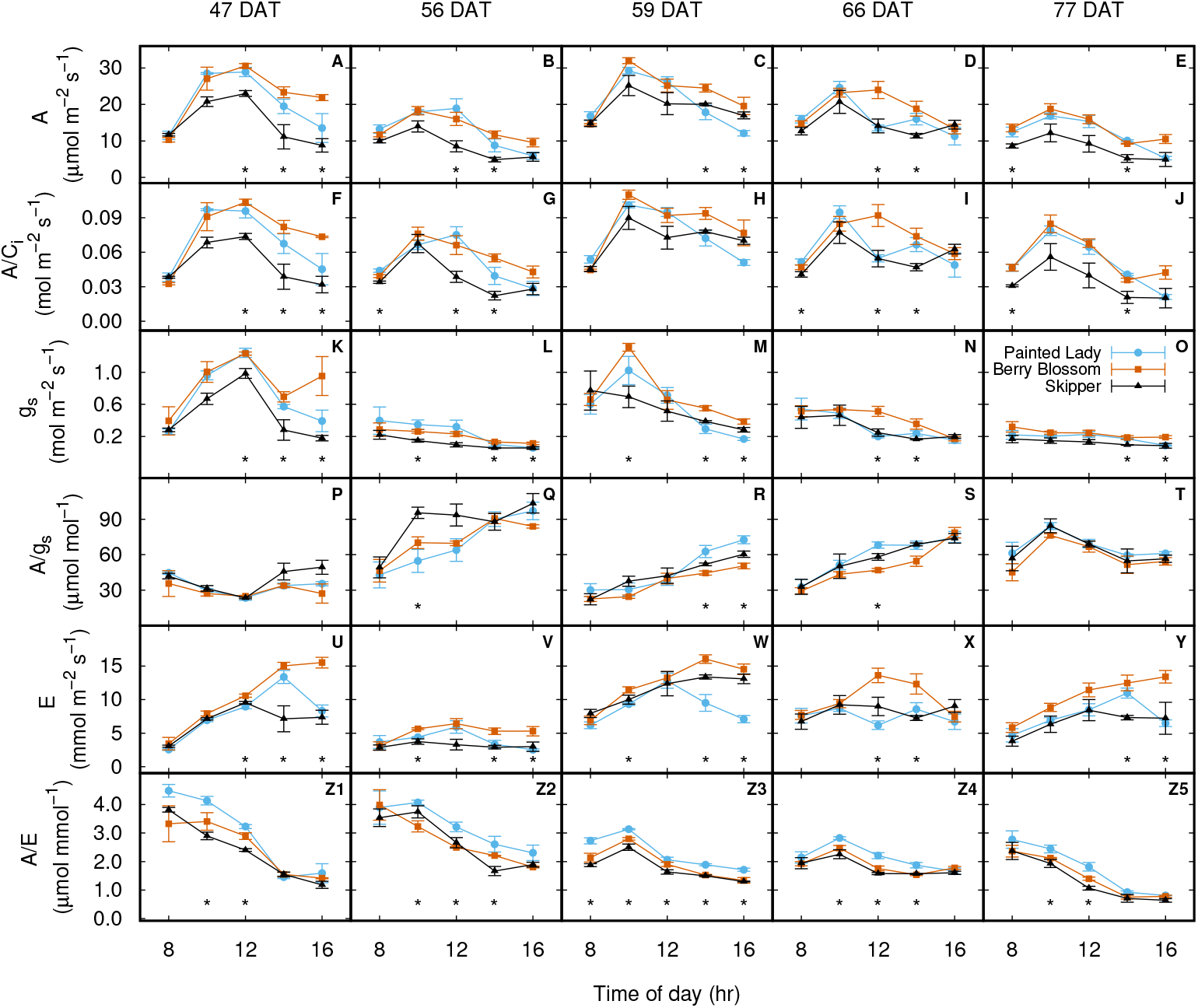
Diurnal courses of net photosynthetic rate (*A* shown in panels **A**-**E**), carboxylation efficiency (*A*/*C*_*i*_, panels **F**-**J**), stomatal conductance (*g*_*s*_, panels **K**-**O**), intrinsic water use efficiency (*A*/*g*_*s*_, panels **P**-**T**), transpiration rate (*E*, **U**-**Y**), and instantaneous water use efficiency (*A*/*E*, panels **Z1** through **Z5**) of three hemp genotypes measured on June 11, 20, 23, 30, and July 11, 2022 (corresponding to 47, 56, 59, 66 and 77 days, respectively, after transplanting, DAT) at Uvalde, Texas. On each day, the measurement was made five times from 8:00 am to 16:00 pm local time, on three randomly chosen sunlit mature leaves from each of the genotypes (*n* = 3, error bars indicate ± one standard errors of the means, same for Figure 3). A star (“*”) placed below the values of a variable measured at a specific time on a specific day indicates the existence of at least one pairwise significant difference in that variable among the three varieties.

**FIGURE 3.**
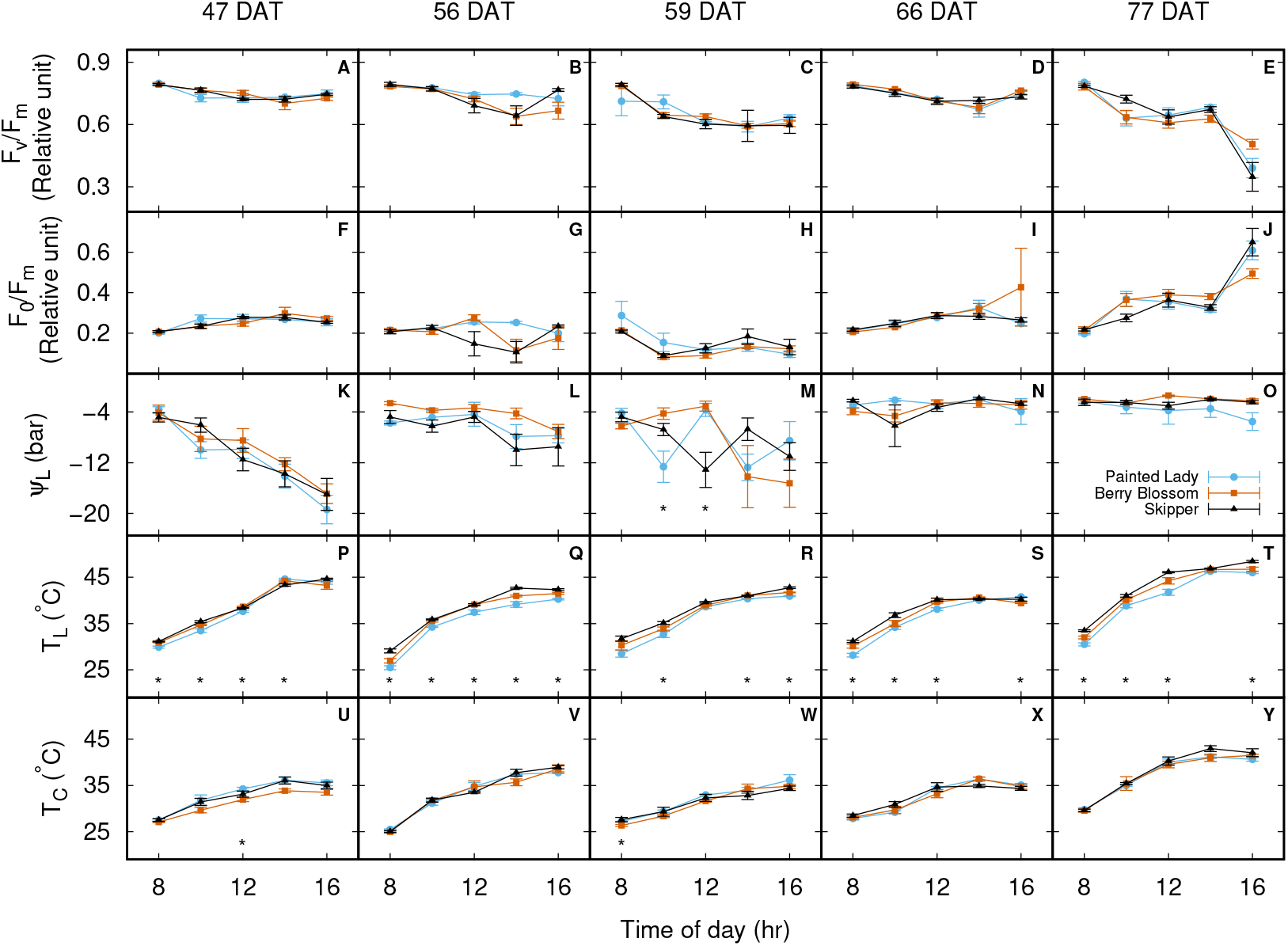
Diurnal courses of photochemical efficiency of PSII (*F*_*v*_ /*F*_*m*_, *n* = 4 leaves; panels **A**-**E**) and thylakoid membrane damage (*F*_*o*_ /*F*_*m*_, panels **F**-**J**), leaf water potential (*ψ*_*L*_, *n* = 4–6, panels **K**-**O**), leaf temperature (*T*_leaf_, *n* = 4, panels **P**-**T**), and canopy temperature (*T*_canopy_, *n* = 4 plants, panels **U**-**Y**) of three hemp varieties measured on June 11, 20, 23, 30, and July 11, 2022 (corresponding to 47, 56, 59, 66 and 77 days, respectively, after transplanting, DAT) at Uvalde, Texas. A star (“*”) placed below the values of a variable measured at a specific time on a specific day indicates the existence of at least one pairwise significant difference in that variable among the three varieties.

The effects of the three-way interactions (*V* × *T* × *D*) show highly dynamical responses of the physiological trait variables of three hemp varieties. However, some salient features in terms of the relationships of leaf photosynthesis and stomatal conductance under various levels of leaf-to-air vapor pressure deficit (*D*) can be identified based on the pooled data of the gas exchange measurements made across the five days. It is evident in Figure 4 that, for all three hemp varieties, *A* (“Photo” or “Photosynthesis”) was highest under high *g*_*s*_ and lower *D*, namely, near the upper-left corner of the contour graphs. That reflects a typical situation in the mid-morning hours of a clear day when the value of *A* was the highest diurnally. The reduction of *A* from the upper-left to the lower-right corner of the graphs approximately corresponded to the trend of change in *A* from mid-morning to the afternoon hours with the simultaneous reduction of *g*_*s*_ and increase of *D* (see Supplementary Figure S2 for the trend of diurnal variations of *D*). We see that, moving from the upper-left to the lower-right corner of the graphs, the largest gradient of *A* occurred in Pained Lady (Figure 4A), when compared with two other varieties. In Berry Blossom, the value of *A* remained high and changed modestly from the morning to the afternoon hours, as indicated by the prevalence of green colors in the contour graph (Figure 4B). Skipper had very low values of *A* in the afternoon hours (as seen in the widespread blue colors in Figure 4C), but in the mid-morning hours, it showed a high value of *A* only briefly, as seen in the restricted dark green area in the upper-left corner of the graph. At 16:00 pm on the warmest day (77 DAT) when *T*_*L*_ was as high as 46.0 to 48.4°C as measured by a thermocouple in the chamber cuvette (Figure 3T), *F*_*v*_ /*F*_*m*_ reached the lowest values (i.e., 0.390, 0.505, 0.348, for Painted Lady, Berry Blossom and Skipper, respectively) and *F*_0_/*F*_*m*_ reached the highest values of the day for all three varieties. The significant reduction in *F*_*v*_ /*F*_*m*_ and increase in *F*_0_/*F*_*m*_ measured at 16:00 pm on 77 DAT also corresponded with an increased *C*_*i*_ /*C*_*a*_ at lower values of *g*_*s*_, especially for Skipper, as seen in Figure 5F, in which the relationship between *C*_*i*_ /*C*_*a*_ and *g*_*s*_ deviated from the generally decreasing trend observed in four other days with a more moderate temperature (Figure 5B-E).

**FIGURE 4.**
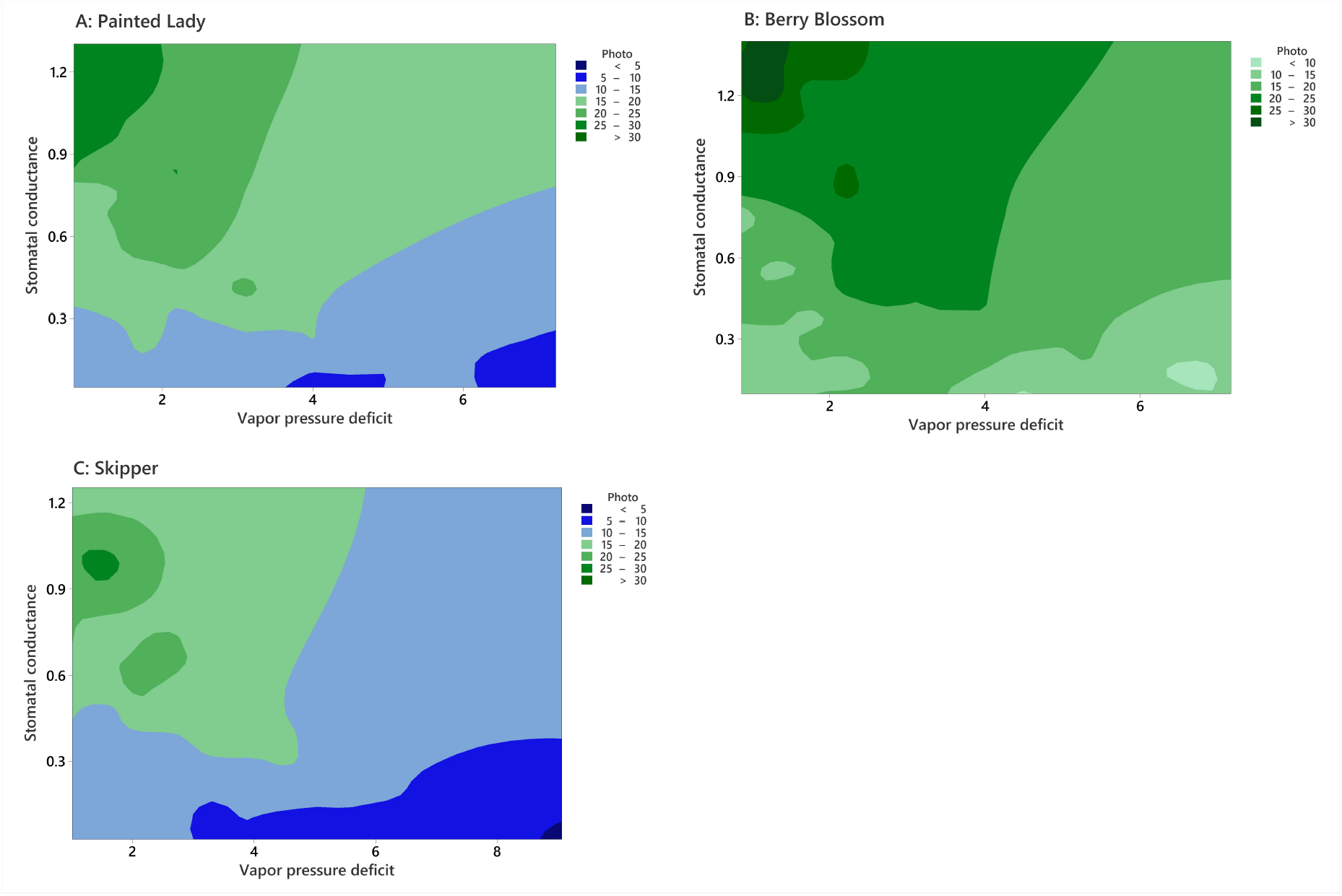
Contour plot depicting net photosynthetic rate (“Photo”) (*µ*mol m^−2^ s^−1^) as a function of stomatal conductance (mol m^−2^ s^−1^) and vapor pressure deficit (kPa) based on leaf gas exchange measurements made on 75 leaves over five clear days for three hemp varieties (same as in Figs. 2 and 3) at Uvalde.

**FIGURE 5.**
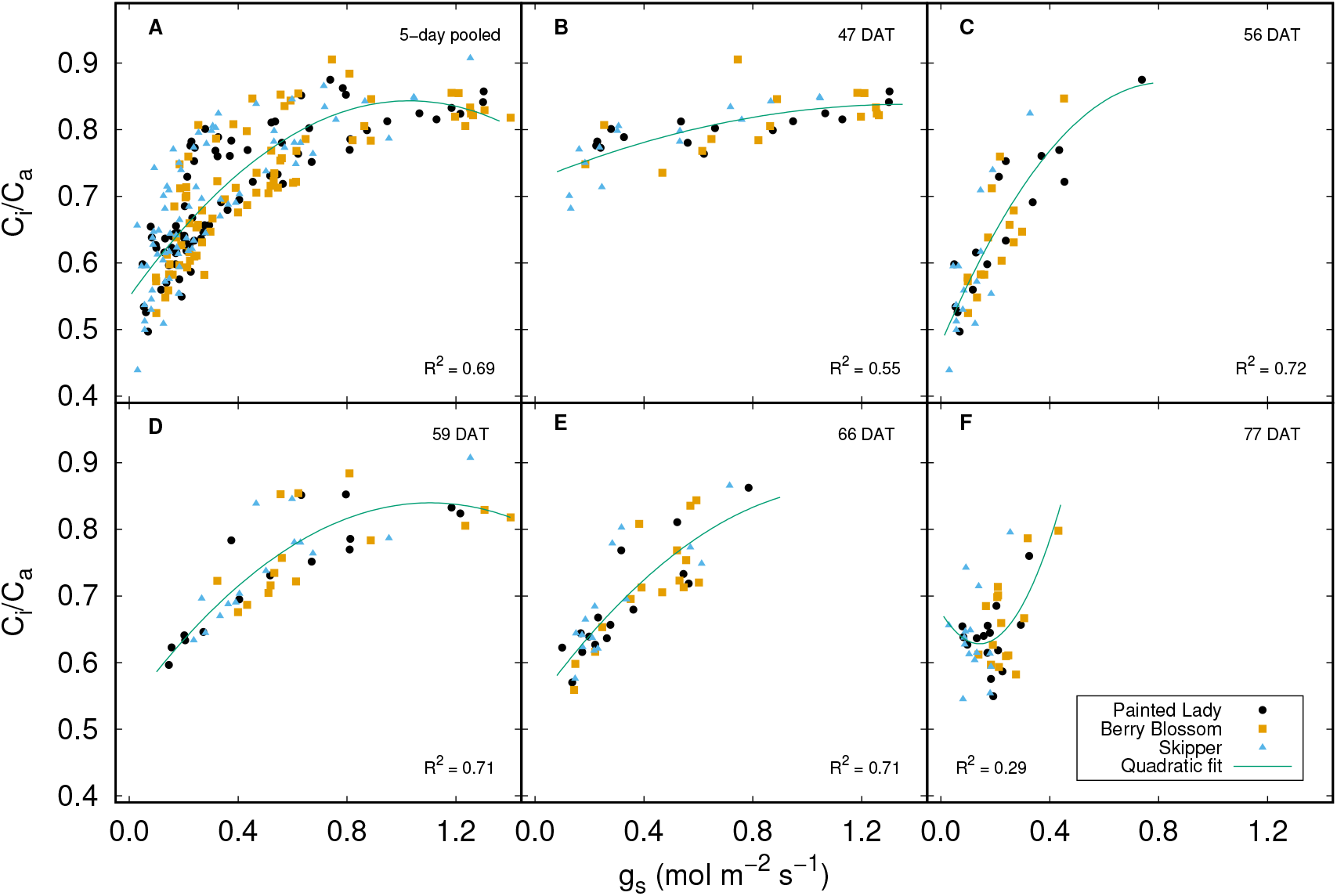
Relationships between the ratios of *C*_*i*_ /*C*_*a*_ and leaf stomatal conductance (*g*_*s*_) for pooled samples collected over five days for three hemp genotypes (panel **A**), and for individual days (panels **B**-**F**). A quadratic function was fit to the data to show a generally decreasing trend of *C*_*i*_ /*C*_*a*_ as *g*_*s*_ decreased, except on 77 DAT (days after transplanting). All regressions are significant (*p* < 0.0001). Each data point represents one leaf.

In Figure 6, almost all data points of *A* of the three hemp varieties are enveloped by a bounding curve described by the quadratic equation 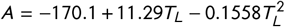, from which the optimal leaf temperature for photosynthesis was estimated as 36.3°C. The highest photosynthetic rate was recorded for Berry Blossom (33.6 *µ*mol CO_2_ m^−2^ s^−1^) at a leaf temperature of 34.6°C at 10:00 am on 59 DAP (see Figure 2C).

**FIGURE 6.**
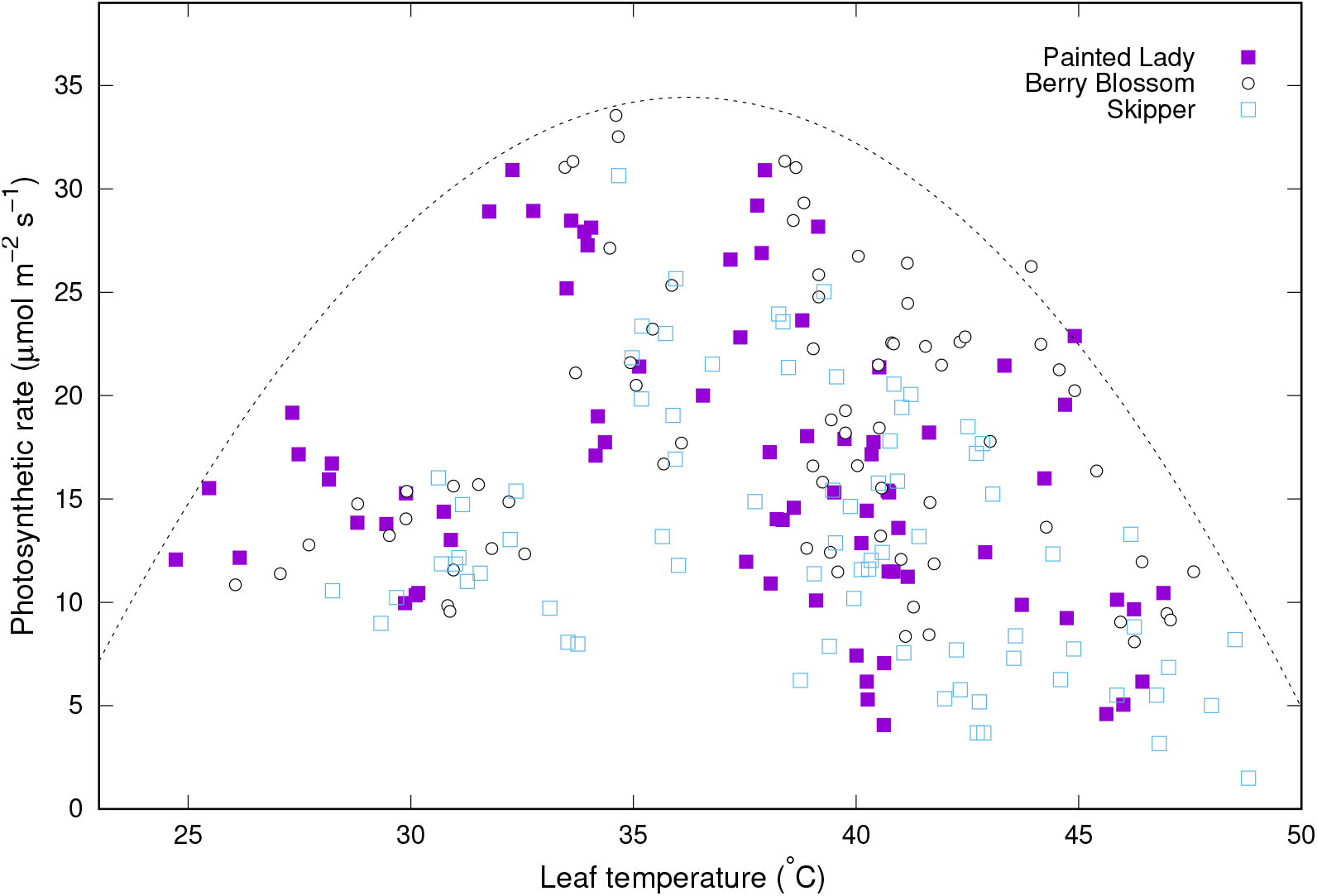
Scatter plot of net photosynthetic rate as a function of leaf temperature for 75 leaves measured each on three hemp genotypes, i.e., Painted Lady, Berry Blossom, and Skipper, on five clear days (June 11, 20, 23, 30, and July 11, 2022) at Uvalde, Texas. On each day, the measurement was made five times from 8:00 am to 4:00 pm local time using a set of LI-6400XT Portable Photosynthesis System. Almost all data points of photosynthetic rate of the hemp genotypes are enveloped by a bounding curve described by the quadratic equation 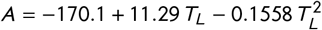, from which the optimal leaf temperature for photosynthesis was estimated to occur at a leaf temperature (*T*_*L*_) of 36.3 °C.

### 3.2 Optimal stomatal behavior through balancing photosynthesis and transpiration in different hemp varieties

The estimated value of parameter *g*_1_ for hemp genotype Painted Lady was significantly lower than the like values for Berry Blossom and Skipper (Table 3). Alternatively, regression of *g*_*s*_ against 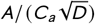 showed that the slope of Painted Lady was significantly lower than those of two other varieties (*P* = 0.033; Figure 7). These results may be considered as an indicator of the overall patterns of the optimal stomatal behavior for the three hemp varieties growing in southwest Texas.

**TABLE 3.** Statistics of fits of the unified stomatal model (Eqn. 1) to the diurnal leaf gas exchange data of three hemp varieties in the Uvalde hemp cultivation trial in 2022.

| Variety | *n* | *g*<sub>1</sub> | *s* |
| --- | --- | --- | --- |
| Painted Lady | 75 | <sup>c</sup>7.58 (0.26) | 0.126 |
| Berry Blossom | 75 | <sup>a</sup>8.87 (0.30) | 0.150 |
| Skipper | 75 | <sup>abc</sup>8.35 (0.34) | 0.121 |

The estimated standard errors for the model parameter are shown in parentheses following the means. For each of the hemp varieties, diurnal gas exchange measurements made on 75 leaves over five days after planting (DAP)(i.e., 47, 56, 59, 66, and 77 DAP) were used. All leaf gas exchange measurements were made on clear days under field conditions. The unit of parameter *g*<sub>1</sub> is kPa<sup>0.5</sup>. The standard error of the regression (*s*) indicates that approximately 95% of the observations should fall within ±2*s* of the predicted *g*<sub>s</sub>.
Different letters indicate statistically significant differences in model parameter values among three hemp varieties (*p* < 0.05).

**FIGURE 7.**
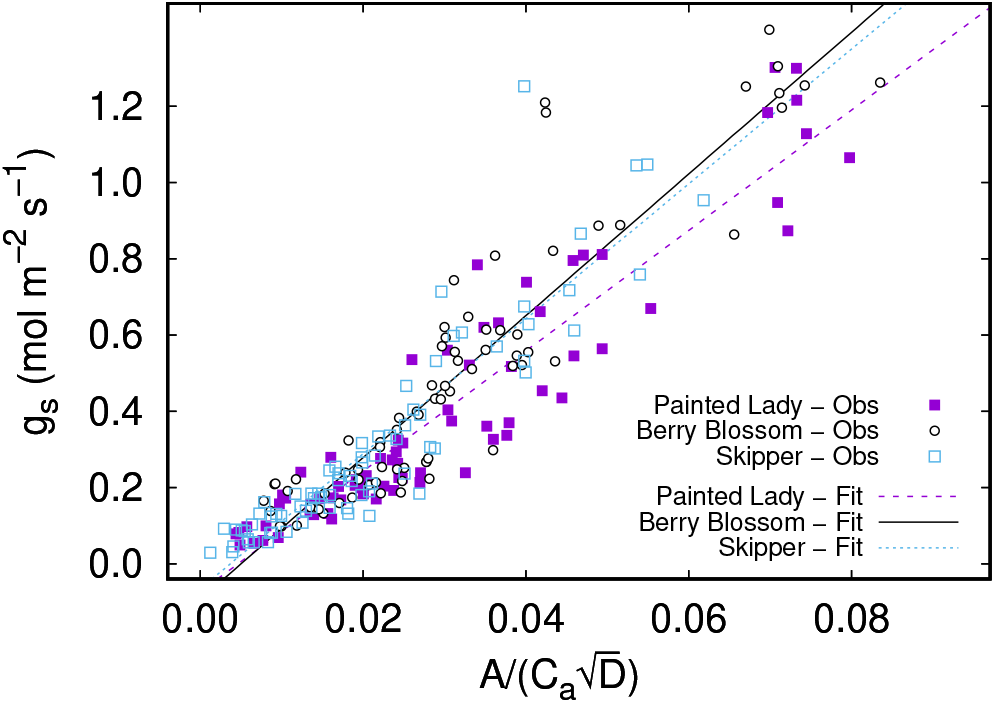
Visualization of stomatal behavior for selected three hemp genotypes by regressing measured values of leaf stomatal conductance (*g*_*s*_, mol m^−2^ s^−1^) against 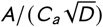, as predicted using the unified stomatal model (Eqn. 1). *A* is net photosynthetic rate (*µ*mol m^−2^ s^−1^), *C*_*a*_ is the atmospheric CO_2_ concentration at the leaf surface (*µ*mol mol^−1^), and *D* is leaf-to-air vapor pressure deficit (kPa). Each of the hemp datasets included gas exchange values of 75 leaves measured five times each day for five days in 2022 (i.e., on June 11, 20, 23, and 30, and July 11, with the diurnal course data showing in Figure 2). All measurements were made on clear days under field conditions. The slopes of the datasets are proportional to the best fits of the model (*g*_1_) as shown in Table 3.

### 3.3 Plant height, biomass accumulation, and root traits of different varieties

The height growth of Painted Lady and Skipper was faster than that of Berry Blossom, with the peak growth rate occurring in May 15 (20 days after transplanting) for Painted Lady (2.9 cm/day) and Skipper (2.3 cm/day) and in May 23 (28 days after transplanting) for Berry Blossom (1.6 cm/day; Figure 1C). A significantly higher plant height was observed in Painted Lady as compared to two other varieties, while the higher inflorescence and shoot dry mass values were observed in Berry Blossom (Table 4). At the same time, however, there was no significant difference in root dry mass or root/shoot ratio (Table 4).

**TABLE 4.** Plant height, biomass accumulation (inflorescence, shoot, and root), root-shoot ratio, and cannabinoid contents of three hemp varieties in the Uvalde hemp cultivation trial in 2022.

|  | Painted Lady | Berry Blossom | Skipper |
| --- | --- | --- | --- |
| Plant height (cm) | <sup>a</sup>115 (4)<sup>†</sup> | 101 (4) | 98.4 (61) |
| Inflorescence mass (kg plant<sup>-1</sup>) | 0.28 (0.06) | <sup>a</sup>0.43 (0.07) | 0.23 (0.002) |
| <sup>‡</sup>Shoot mass (kg plant<sup>-1</sup>) | 0.32 (0.07) | <sup>a</sup>0.48 (0.07) | 0.26 (0.03) |
| <sup>#</sup>Root mass (kg m<sup>-3</sup>) | 0.13 (0.009) | 0.15 (0.02) | 0.12 (0.007) |
| Root/Shoot | 0.54 (0.18) | 0.34 (0.04) | 0.47 (0.04) |
| Stem basal diameter (mm) | 21.5 (1.1) | 20.2 (1.4) | 19.7 (0.3) |
| <sup>¶</sup>Cannabinoid contents |  |  |  |
| <sup>§</sup>CBD (mg g<sup>-1</sup>) | 43.3 (2.2) | 25.0 (0.84) | 50.0 (4.89) |
| THC (mg g<sup>-1</sup>) | 0.88 (0.16) | 0.69 (0.08) | 1.09 (0.10) |
| THC (%) | 0.09 (0.008) | 0.07 (0.002) | 0.11 (0.008) |
| THC/CBD | 0.020 (0.004) | 0.028 (0.003) | 0.022 (0.003) |

- <sup>†</sup> Data shown are means followed by standard errors of the means (SEM) in parentheses. The average value marked by letter "a" is significantly higher than the mean value in that row ( $\alpha = 0.10$ ).
- <sup>§</sup> CBD, cannabidiol; THC, Δ-9-tetrahydrocannabinol.
- <sup>‡</sup> Values of the shoot and root mass are averages of plants from each variety from the five blocks ( $n = 5$ ). For each variety, the value for each block is the average of four plants ( $n = 4$ ) grown in that block.
- <sup>#</sup> Root mass (kg m<sup>-3</sup>) is defined as estimated total root mass (fine roots + thick roots) found in 1 m<sup>3</sup> of soil volume within 1 m soil depth profile, assuming for top 10-cm of soil the thick roots only exists in the 25 cm diameter area centering the plant.

The allometric relationships between pairs of selected biomass and morphological variables are established and the results are shown in Figure 8 and Table 5. Regardless of varieties, there was a highly strong linear relationship between log-transformed shoot dry mass and shoot fresh mass (Figure 8A) and log-transformed inflorescence dry mass and inflorescence fresh mass (Figure 8B). The relationship between log-transformed inflorescence dry mass and stem dry mass was also strong (Figure 8D) with an *R*^2^ of 0.860 (Table 5), while the relationship between log-transform root dry mas and shoot dry mass was relatively weak (Figure 8C; Table 5). When using measured basal diameter of individual plants to predict the dry mass of shoot or inflorescence, different equations were established for three hemp varieties (Figure 8E-F), with Skipper having the lowest Y-intercept and Painted Lady the highest (Table 5).

**TABLE 5.** Further details of the allometric equations shown in Fig. 8.

| Relationship | Variety | ln (b) | k | n | R<sup>2</sup> | P-value |
| --- | --- | --- | --- | --- | --- | --- |
| ln (shoot-dry mass) - ln (shoot-fresh mass) | † Combined | -1.127 | 1.007 | 59 | 0.996 | < 0.0005 |
| ln (infl.-dry mass) - ln (infl.-fresh mass) | Combined | -1.184 | 1.102 | 59 | 0.995 | < 0.0005 |
| ln (root-dry mass) - ln (shoot-fresh mass) | Combined | 3.503 | 0.3321 | 58 | 0.384 | < 0.0005 |
| ln (infl.-dry mass) - ln (stem-dry mass) | Combined | 1.692 | 1.002 | 60 | 0.860 | < 0.0005 |
| ln (infl.-dry mass) - ln (stem basal diameter) | Painted Lady | 0.496 | 1.590 | 18 | 0.527 | 0.001 |
|  | Berry Blossom | 0.0659 | 1.828 | 18 | 0.652 | < 0.0005 |
|  | Skipper | -4.633 | 3.253 | 19 | 0.578 | < 0.0005 |
| ln (shoot-dry mass) - ln (stem basal diameter) | Painted Lady | 0.8150 | 1.543 | 18 | 0.541 | < 0.0005 |
|  | Berry Blossom | 0.3579 | 1.788 | 18 | 0.648 | < 0.0005 |
|  | Skipper | -4.407 | 3.233 | 19 | 0.590 | < 0.0005 |

† Data of the three varieties were combined if the slopes and y-intercepts were not significantly different ( $P > 0.05$ ). For a specific allometric relationship, different letters indicate significant differences in parameters among varieties ( $P < 0.054$ ).

**FIGURE 8.**
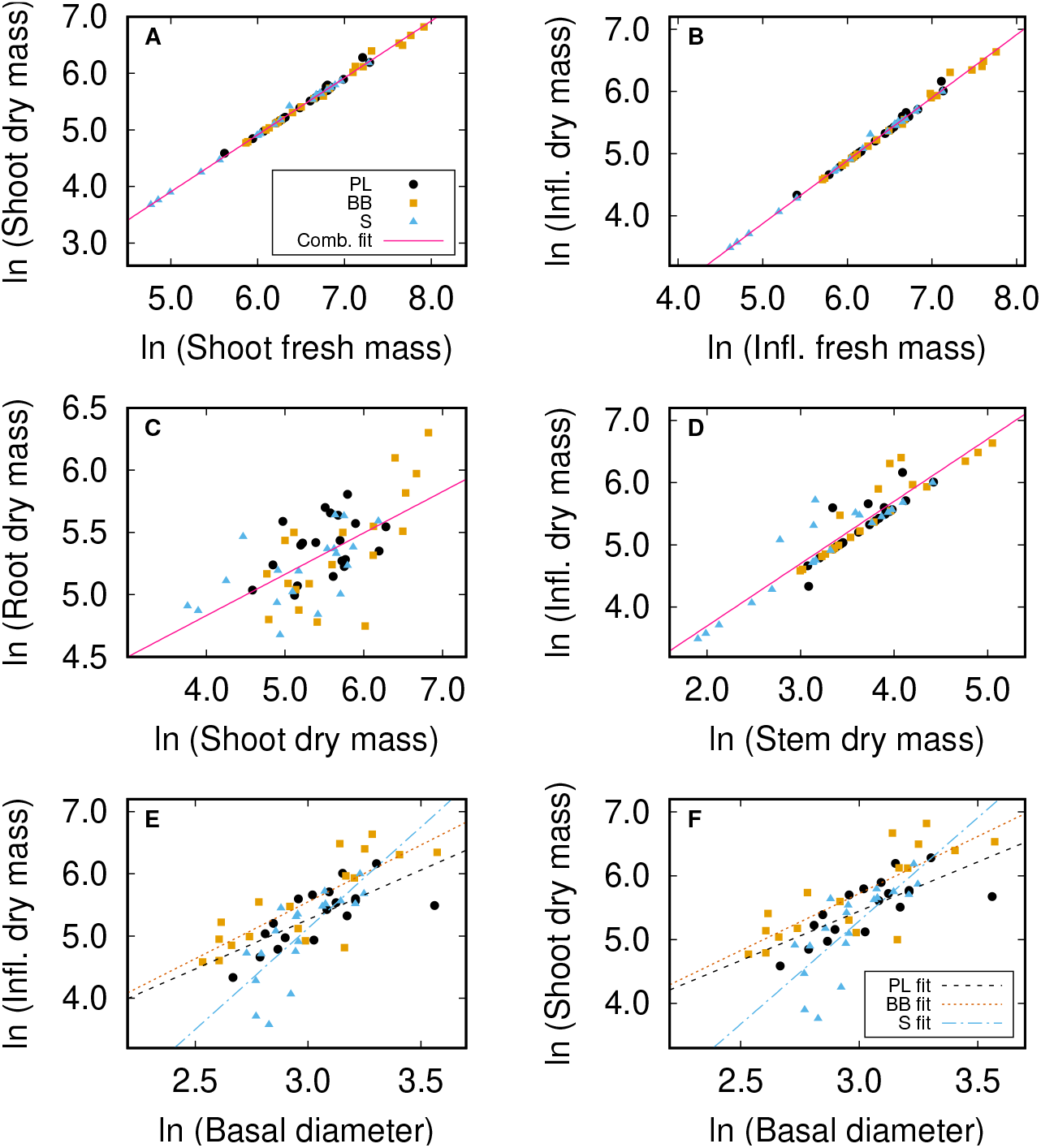
Linear regression(s) of ln-transformed (**A**) shoot dry mass and shoot fresh mass, (**B**) inflorescence dry mass and inflorescence fresh mass, (**C**) root dry mass and shoot dry mass, (**D**) inflorescence dry mass and stem dry mass, (**E**) inflorescence dry mass and basal diameter, and (**F**) shoot dry mass and basal diameter for three hemp varieties at Uvalde in 2022. All the regression equations share the form of *y* = *bx*^*k*^, where the unit of biomass is g plant^−1^ and that for basal diameter is mm. If there was no difference in slope or the y-intercept among the varieties, a common regression line is shown (Panels **A**-**D**); otherwise, the individual regression lines for the three varieties are shown (Panels **E**-**F**). The coefficients of the equations and their comparisons are presented in Table 5. Abbreviations: Comb. = Combined; Infl. = Inflorescence; PL = Painted Lady; BB = Berry Blossom; S = Skipper.

Four of the eight root trait/ root distribution parameters showed significant differences among the hemp varieties. Of the three varieties, Skipper had the highest value of specific root length (SRL) and Berry Blossom had the shallowest rooting depth, as seen in the high values of *α*_*r*_ and *β*, and low value of *Z*_95_ (Table 6). The root distribution patterns of the three hemp varieties were visualized by plotting the measured and fitted root length density as a function of soil depth (Figure 9).

**TABLE 6.** Selected root traits and parameters of root distribution for three hemp varieties grown at Uvalde in 2022 under field conditions.

| Variety | SRL (cm mg<sup>-1</sup>) | D (mm) | z<sub>95</sub> (cm) | R<sub>t2b</sub> | K | α<sub>r</sub> | β |
| --- | --- | --- | --- | --- | --- | --- | --- |
| PL | 10.96 (0.30) | 0.35 (0.01) | 74.2 (5.05) | 3.11 (0.41) | 3.04 (0.41) | 0.028 (0.004) | 2.38 (0.40) |
| BB | 11.00 (0.60) | 0.37 (0.01) | <sup>b</sup>63.0 (4.37) | 3.16 (0.19) | 4.62 (0.89) | <sup>a</sup>0.046 (0.009) | <sup>a</sup>3.54 (0.59) |
| S | <sup>a</sup>12.31 (0.42) | 0.35 (0.01) | 77.3 (2.92) | 2.62 (0.20) | 3.12 (0.38) | 0.027 (0.003) | 2.17 (0.25) |

- Abbreviations: **PL**, Painted Lady; **BB**, Berry Blossom; **S**, Skipper. - Definition of variables: **SRL**, specific root length for fine roots; **D**, average diameter of fine roots; z<sub>95</sub>, rooting depth defined as soil depth above which 95% of fine roots is located; R<sub>t2b</sub>, ratio of total root length between top and bottom soil layers (divided at 40 cm depth); K, the potential root length density near soil surface; α<sub>r</sub> and β, the rate of reduction of root length density with the increase of soil depth. Parameters z<sub>95</sub>, R<sub>t2b</sub>, K, α<sub>r</sub>, and β were calculated using the computer program of Dong et al. (2019). - Data shown are means followed by standard errors of the means (SEM) in parentheses. - Within a column, capital letter "a" and "b" indicate a significantly higher or lower value as compared with the mean value (α = 0.10, n = 5).

**FIGURE 9.**
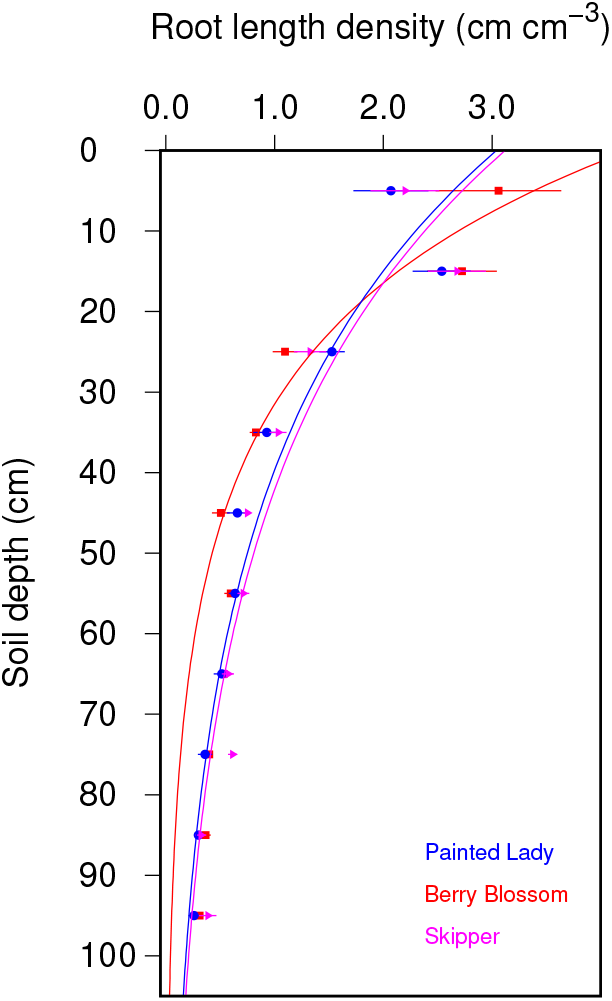
Vertical distributions of root length density (defined as the length of total fine roots found at 1 cm^3^ of soil) for the three hemp varieties measured at Uvalde near end of the 2022 growing season. Each data point represents the average of 5 values of root length density (for 5 plots) of a hemp variety measured at a specific soil depth interval. Error bars indicate ± 1 standard errors of means (*n* = 5). The distribution of root length density (*L*_*r*_) as a function of soil depth (*z*) was described by equation 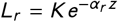 (Dong et al., 2019), *where the values of K* and *α*_*r*_ were obtained from Table 6.

### 3.4 Cannabinoid profiling of different genotypes

The occurrence of the peak flowering stage (at BBCH scale 65-67) was similar in Painted Lady and Skipper, with the highest frequency being recorded on July 5 and July 7 respectively (Supplementary Figure S2). For Berry Blossom, however, the occurrence of the peak flowering stage spanned a wider time frame, with the highest frequency taking place on July 23, 2022. Although the total CBD and THC values of Skipper were numerically higher than those of two other varieties, no statistically significant difference was observed from our data (Table 4). The three varieties had similar THC/CBD ratios, which are significantly lower than those reported for selected hemp accessions from India (Khajuria et al., 2020). In one of the five blocks, the measured values of CBDA for Painted Lady and Skipper were very high. However, no statistically significant difference was found in CBDA among the three hemp varieties (see Supplementary Figure S3). Overall, the %THC values across the three hemp varieties were below 0.3%.

## 4 DISCUSSION

Our field data suggest that the tested three hemp varieties exhibited interesting adaptations to the higher temperatures of southwest Texas under well-irrigated conditions, with the optimal temperature for photosynthesis being 1.3 °C higher (see Figure 6) than the results obtained for hemp from contrasting environments(Bazzaz et al., 1975; Chandra et al., 2011; Cosentino et al., 2013; Tang et al., 2017). The highest net photosynthetic rate (*A*) measured in our study (Figure 6) was higher than the like values reported for hemp in several European studies (Tang et al., 2017, 2018; Herppich et al., 2020). There were 81 out of 159 days in the 2022 growing season at Uvalde (Figure 1A) during which the maximum daily air temperature was greater than the optimal temperature of photosynthesis (i.e., 36.3 °C). Also note that July 2022 was ranked among the historically top 5 warmest to the warmest in southwest Texas since 1895 (see Supplementary Figure S5). Yet photosynthesis of the three hemp varieties was dominated by stomatal limitation (Figure 5B-E, see also Brodribb (1996) for examples in some coniferous species), except in the late afternoon of a very warm day (77 DAT) when non-stomatal inhibition, such as thylakoid membrane damage and reductions in both PSII efficiency and overall net photosynthetic rate showed up (see Figure 2E; Figure 3E,J). Diurnal trends of *F*_*v*_ /*F*_*m*_ and *F*_0_/*F*_*m*_ measured in hemp in our study were similar to those of cotton measured at the same site in 2022 (Dong et al., 2023). Although the values of *Fv /Fm* tended to reduce during the daytime hours when the chlorophyll fluorescence was measured, the starting values at 8:00 am on the five days for all three varieties were higher than 0.78, except in Painted Lady on 59 DAT when the starting value of *F*_*v*_ /*F*_*m*_ was 0.712 (Figure 3C). Since *F*_*v*_ /*F*_*m*_ values at or higher than 0.78 represent typical values of unstressed healthy plants (Björkman and Demmig, 1987; Brodribb, 1996; Guidi et al., 2019; Li et al., 2019), the recovery of high *Fv /Fm* values at 8:00 am in our data suggest that the possible photochemical inhibition of photosynthesis as indicated in the diurnal reduction in *F*_*v*_ /*F*_*m*_ was probably transient in nature, a result presumably related to the acclimation of the photosynthetic apparatus of the hemp varieties to high temperatures of southwest Texas, similar to the photosynthetic acclimation to high temperatures in maize (Sinsawat et al., 2004). As a matter of fact, the *F*_*v*_ /*F*_*m*_ values in early morning hours in our study were generally higher than the value for hemp (≈ 0.74-0.75) reported by Mejía-Londoño et al. (2023). As a result, overall, the naturally occurring high temperatures at Uvalde did not significantly affect the photosynthetic parameters of the tested three hemp varieties in 2022 (our Question #1).

To address our second question, the effect of hemp’s deep-root traits did not show clear evidence of affecting hemp physiology in our test. Although the rooting depth of Berry Blossom was shallower than that of two other varieties (Table 6 and Figure 9), its photosynthesis generally was higher than either of two other varieties (Figure 2A-E). One possible explanation is that the hemp varieties were under irrigated production and there was no severe water stress developed, as seen from the relatively less negative leaf water potential (*ψ*_*L*_, Figure 3K-O) and slight differences in leaf and canopy temperatures among three varieties (Figure 3P-Y) in the five clear days. The total root mass of three hemp varieties in our study varied from 0.12 to 0.15 kg m^−2^, or 1.2 t ha^−1^ to 1.5 t ha^−2^, which was about 50% of the values reported in an Italian study (Amaducci et al., 2008). Likewise, the maximum root length density (RLD) at the top 10 cm soil layer in our study was slightly more than 50% of the value reported in Amaducci et al. (2008). But the root/shoot ratio in our study (0.34-0.54) was more than twice that of the Italian study, indicating a much lower shoot mass in our hemp plants.

Nor did we observe significant differences in cannabinoid contents in relation to environmental and varietal responses, although the small-statured Skipper seemed to have numerically higher cannabinoid contents than two other varieties. Khajuria et al. (2020) used both the hemp accessions with a wide range of THC levels and *Arabidopsis thaliana* treated with varying doses of THC to demonstrate that a high THC level caused reductions in photochemical efficiency and the non-photochemical quenching (NPQ) mechanism that help to dissipate excitation energy and maintain the integrity of photosystem II (PS II). Some of their high THC accessions had a THC content > 6%. In the current study, there was no clear relationship between the measured photochemical efficiency and the THC content. It is possible that the THC levels in our hemp varieties might be too low to cause a negative impact on the photosynthesis machinery. This needs to be confirmed by further studies. The observed deep-rooting tendency in Skipper and Painted Lady may potentially give these varieties an advantage in water-limited conditions, as has been found in sugar beet (Bodner and Alsalem, 2023). This also needs to be confirmed in future studies.

Our leaf photosynthesis data offered an opportunity for a deeper understanding of the field adaptations and potential water use efficiency of the hemp varieties. This was approached through the stomatal optimization model of Medlyn et al. (2011), which synthesized the key ideas of prior models (Ball et al., 1987; Leuning, 1995, e.g.), and has been applied in diverse plant species (Lin et al., 2015; Gardner et al., 2023). However, the stomatal optimization model has rarely been used to directly interpret the diurnal leaf gas exchange data under field conditions. When discussing the differences of parameter *g*_1_ (see Eqn 1 in Materials and Methods for definition) among different plant species of diverse taxonomy, Medlyn et al. (2011) used the term ‘conservatory’ to describe a stomatal strategy that tends to curb excessive water loss while allowing CO_2_ diffusion into the leaf mesophyll cells for photosynthesis to occur. However, it remains unknown how this ‘conservatory’ stomatal strategy can be applied to interpret the diurnal changes of net photosynthesis (*A*), stomatal conductance (*g*_*s*_) and vapor pressure deficit (*D*). The relationships among these variables can be illustrated using the diurnal-course photosynthesis data of our hemp varieties.

Our measured leaf gas exchange data suggested that Painted Lady displayed a more efficient strategy of water use, compared with two other hemp varieties (Table 3 and Figure 7). This efficient strategy, although not being reflected in the intrinsic water use efficiency (*A*/*g*_*s*_), was present in the daily instantaneous water use efficiency (*A*/*E*) (Table 2; Figure 2Z1-Z5). Furthermore, this is demonstrated in the ways in which *g*_*s*_ and *A* varied diurnally: Painted Lady’s stomatal behavior was more efficient for photosynthetic carbon gain because it tended to fully use the morning hours (when *D* was lower; see Supplementary Table S2 for values of *D* in the five measurement days) to do photosynthesis with opened stomata, while reducing photosynthesis with limited stomatal opening in the afternoon hours when *D* was high (see blue lines in Figure 2A-E, K-O;Figure 4A). On the other hand, the stomatal strategy of Berry Blossom was not as efficient as that of Painted Lady because, while it also fully used the morning hours to do photosynthesis with large stomatal conductance (similar as Painted Lady), it tended not to strongly curb photosynthesis (and water loss) in the afternoon hours when *D* was high (see the orange lines in Figure 2A-E, K-O and also Figure 4B). Lastly, the stomatal strategy of Skipper was less efficient because the diurnal values of both *A* and *g*_*s*_ tended to be the lowest among the three varieties across five days (see the black lines in Figure 2A-E, K-O and and Figure 4C), meaning that it was inclined to miss the ‘golden’ morning hours to increase photosynthesis. The above trend is summarized also in Figure 4.

Although Berry Blossom had a higher photosynthetic rate, a higher shoot biomass and inflorescence biomass compared with two other hemp varieties tested in this study, it yielded similar CBD production as compared to two other varieties (Table 4). Further study in this direction is needed to test the regional adaptation of these hemp varieties in relation to the CBD yield. Our allometric analysis on biomass and morphological traits indicate that there are opportunities for using easily measured traits to predict or estimate traits that are more time-consuming or more difficult to measure under field conditions. In particular, the process of drying the freshly harvested hemp inflorescent takes a long time under a low temperature condition to prevent terpenes from being lost through evaporation. In this case, the inflorescent dry mass may be estimated from the fresh mass using the allometric equations similar to the one reported in this work. Also, as it is typically time-consuming to estimate root biomass and rooting depth of hemp under field conditions, equations similar as the ones developed in this work may be used to get a first estimate of root dry mass, even though the strength of the regression relationship is modest (e.g., Figure 8C).

To conclude, the three hemp varieties displayed varied behavior strategies in southwest Texas through their distinct stomatal responses and growth habits. Their photosynthetic parameters and cannabinoid contents remained largely unaffected by the high summer temperatures of the region (in comparison to the optimal temperature for photosynthesis of hemp). Painted Lady exhibited a fast-growing, deep-rooting habit and a potentially more efficient stomatal strategy to strive in water-limited conditions. Skipper, despite having a lower shoot biomass, also exhibited a fast-growing and deep-rooting habit, and slightly higher cannabinoid contents in its inflorescence. Berry Blossom matured later than the two other varieties, but it showed a high shoot biomass accumulation. It may also potentially use more water by having a high leaf gas exchange rate and an extended length of growth period, which needs to be ascertained in future research. Rigorous studies regarding impacts of environmental stresses on cannabinoids are limited, and those that exist—some under greenhouse conditions (Khajuria et al., 2020), others in field conditions but only short-term stress (Toth et al., 2021)—are conflicting in results. Additional research in this direction is needed to take into consideration the genetic origin and its tendency toward CBD vs. THC biosynthesis.

## Author contributions

X.D., R.W.J., and J.S.V.S. conceived and designed the experiment; D.D.B., R.W.J., X.D. and D.I.L. secured funding; J.S.V.S., X.D. and M.V.J. performed the experiment; X.D., J.S.V.S., R.W.J., D.D.B. and D.I.L. analyzed and interpreted the data; J.S.V.S., X. D. and M.V.J. wrote the paper with inputs and approval from all authors.

## Acknowledgments

The funding of this work came from “Hemp Cultivation and Cannabinoid Production,” which was provided by Rare Earth Genomics Texas, LLC. to R.W.J., “SUB to AL-RSCH* Commercial Analysis of HEMP,” provided by Green Ocean Sciences, Inc. DBA Ionization Labs to D.D.B., as well as funding from the USDA NIFA Hatch/Multistate projects to X.D., R.W.J., D.I.L and D.D.B. We thank staff members at the Uvalde Research Center for help in transplanting, and especially we appreciate Dean Hillis for providing crop management, Benjamin Puerta for technical support, Carrie Hensarling for measuring plant nutrient concentration, and Christine Thompson and Liza Silva Sanchez for administrative support. We especially appreciate Mark Hernandez and Joe Moreno at Southwest Texas Junior College for assistance in field and laboratory measurements. Finally, we thank Janet Patton, Jennifer and Jessica Dong for editing the manuscript.

## Supporting Information

There are ten supplementary files associated with this article. Supplementary Figure S1 shows the appearance of plants as growing in the field conditions and a few measurement scenes. Supplementary Figure S2 shows the diurnal variation of solar radiation, air temperature, relative humidity, vapor pressure deficit, and wind speed during the five days in 2022 during which the leaf gas exchange of the three hemp varieties was measured. Supplementary Figure S3 provides a visualization of the occurrences of the peak flowering stage (BBCH scale 65-67) of the three hemp varieties. Supplementary Figure S4 shows the variability of the individual cannabinoid species measured for the three hemp varieties at the peak flowering stage. Supplementary Figure S5 highlights July 2022 as one the warmest in the past century in the study region. Supplementary Tables S1-S5 provide detailed comparisons of the variety (*V*) × time of day (*T*) interactions for selected physiological traits of the hemp varieties measured on five clear days in 2022.

Additional data in support of this article, including the gnuplot scripts to reproduce most of the figures, are available at https://doi.org/10.5281/zenodo.10951516.

